# Modeling the Neurocognitive Dynamics of Language across the Lifespan

**DOI:** 10.1101/2023.07.04.547510

**Authors:** C. Guichet, S. Banjac, S. Achard, M. Mermillod, M. Baciu

## Abstract

Healthy aging is associated with a heterogeneous decline across cognitive functions, typically observed between language comprehension and language production (LP). Examining resting-state fMRI and neuropsychological data from 628 healthy adults (age 18-88) from the CamCAN cohort, we performed state-of-the-art graph theoretical analysis to uncover the neural mechanisms underlying this variability. At the cognitive level, our findings suggest that LP is not an isolated function but is modulated throughout the lifespan by the extent of inter-cognitive synergy between semantic and domain-general processes. At the cerebral level, we show that DMN (Default Mode Network) suppression coupled with FPN (Fronto-Parietal Network) integration is the way for the brain to compensate for the effects of dedifferentiation at a minimal cost, efficiently mitigating the age-related decline in LP. Relatedly, reduced DMN suppression in midlife could compromise the ability to manage the cost of FPN integration. This may prompt older adults to adopt a more cost-efficient compensatory strategy that maintains global homeostasis at the expense of LP performances. Taken together, we propose that midlife represents a critical neurocognitive juncture that signifies the onset of LP decline, as older adults gradually lose control over semantic representations. We summarize our findings in a novel SENECA model (Synergistic, Economical, Nonlinear, Emergent, Cognitive Aging), integrating connectomic and cognitive dimensions within a complex system perspective.

**Highlights:**

- Lexical production (*LP*) relies on the interplay between domain-general and semantic processes throughout life.
- DMN (*Default Mode Network*) suppression cooperates with FPN (*Fronto-Parietal Network*) integration to maintain LP performance at a minimal cost.
- Midlife marks a neurocognitive shift, with reduced DMN suppression prompting a more cost-efficient compensatory strategy that prioritizes homeostasis over LP performance.

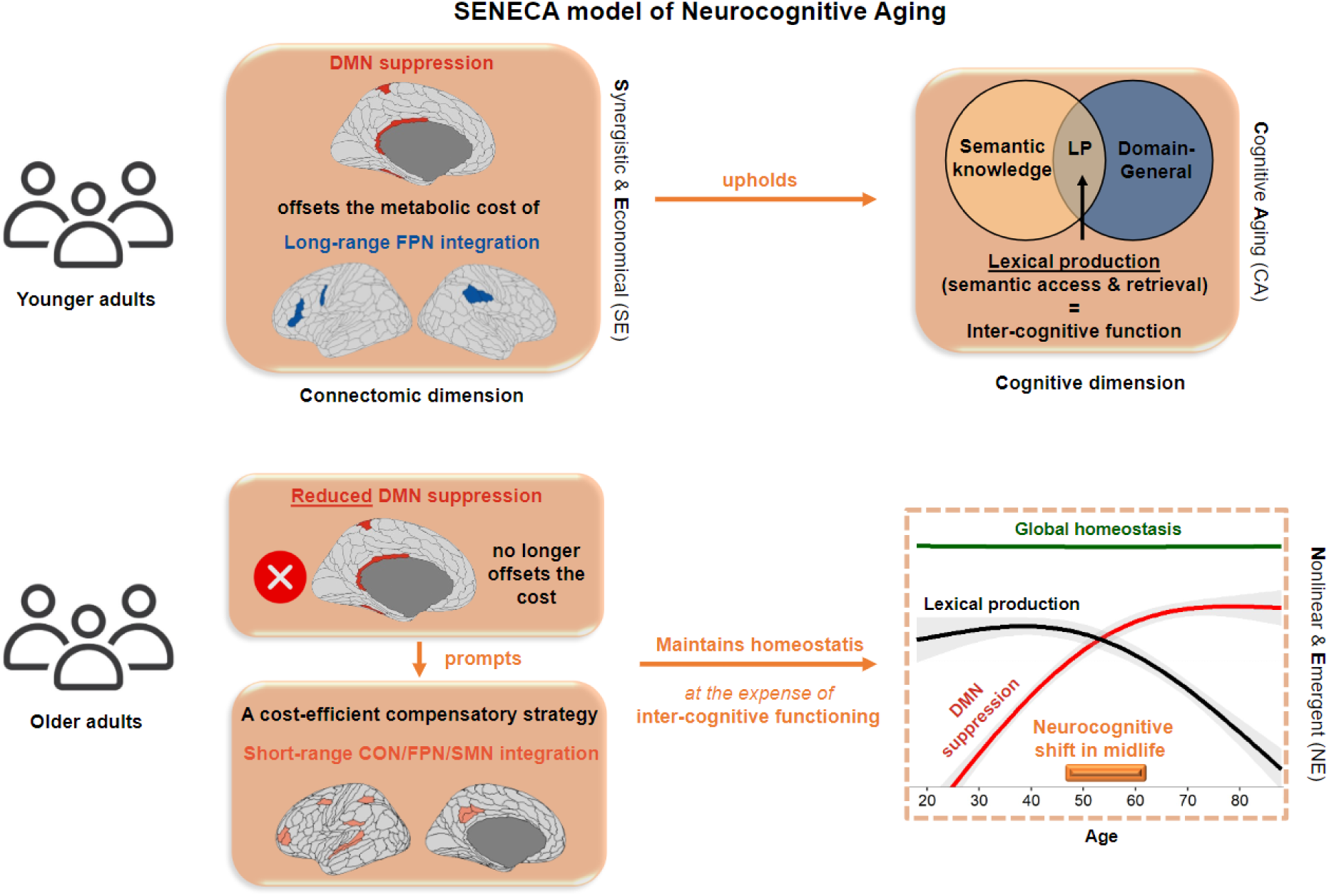

## 1. Introduction

The crucial need to comprehend the mechanisms that uphold normal cognitive functioning during aging arises from the significant increase in the proportion of older individuals (United Nations, 2023). A gradual decline in cognitive abilities typically accompanies aging. However, this decline is heterogeneous across cognitive functions, and some of them, such as language, semantic memory, and vocabulary, tend to be preserved longer (Loaiza, 2024). Therefore, this study aims to shed light on the neural mechanisms that underpin this variability, jointly exploring the functional brain architecture at rest and language-related performance. Specifically, we hypothesize that changes in language function across the lifespan are subserved by the reorganization of the language neurocognitive architecture within the framework of an inter-cognitive interaction between language, long-term memory, and executive functions.

From a cognitive standpoint, language decline under the effect of age is not uniform (Baciu et al., 2021). Although overall language performances tend to be preserved, some linguistic operations may be impaired with age (Ramscar et al., 2014; Wlotko et al., 2010). Indeed, while language comprehension (LC) demonstrates higher resilience to the effects of aging (Diaz et al., 2016; Rossi & Diaz, 2016), language production (LP), which involves lexical retrieval and generation (Baciu et al., 2016, 2021), tends to be more significantly impaired with age (Evrard, 2002; Ramscar et al., 2014). This is particularly obvious in tip-of-the-tongue situations, where individuals experience knowing the meaning of a word but struggle to recall and produce the word form (Burke et al., 1991; Condret-Santi et al., 2013). This discrepancy between LC and LP during aging can be attributed to the advantages of semantic context and the accumulation of semantic knowledge throughout the lifespan (Jongman & Federmeier, 2022; Salthouse, 2019).

From a cerebral perspective, brain networks interact with one another, reflecting how language adapts to socio-communicative contexts by drawing memories, knowledge, and beliefs from long-term memory (Duff & Brown-Schmidt, 2012; Horton, 2007) under the control of executive functions (Corballis, 2019; Hertrich et al., 2020). While long-term memory provides ‘traffic’ content and coherency, executive functioning provides top-down flexibility and coordination to focus, plan, accomplish tasks, and control emotions. In a previous teamwork (Roger, Banjac, et al., 2022), our team proposed a theoretical framework that conceptualizes the inter-cognitive synergy between language, long-term memory, and executive functions at the cognitive level, L∪M (i.e., Language/union/Memory), and suggested that functional connectivity-based interactions may implement this synergy at the neural level.

Indeed, a data-driven analysis highlighted that the language neurocognitive architecture based on extrinsic brain activity (Roger, Rodrigues De Almeida, et al., 2022) comprises four spatially non-overlapping subsystems, each probabilistically mapping onto known resting-state brain networks (i.e., RSNs; Ji et al., 2019): core Language (Net1), Control-Executive (Net2), Conceptual (Net3), and Sensorimotor (Net4). Interestingly, these findings indicate that age-related decline in language production impacts extra-linguistic components (Net2 and Net3) beyond the typical core language network (Hertrich et al., 2020). This suggests that language performances in older adults could be determined by synergistic processing (Gatica et al., 2021), that is the cooperation between control-executive and conceptual/associative processes. In line with (Luppi et al., 2024), we refer to synergistic processing as the joint information that exceeds the sum of each subsystem’s functional connectivity changes. In light of the effect that the reorganization of the language connectome has on language function, we also proposed adopting a ‘cognitomic’ perspective (Roger et al., 2018), emphasizing the constraints that connectomic architecture places on cognitive performances across the lifespan.

Within this perspective, graph theory (Bullmore & Sporns, 2009; Bullmore & Bassett, 2011; Rubinov & Sporns, 2010) is appropriate to describe the connectomic underpinnings of language. Specifically, the brain network can be characterized in terms of integration and segregation properties (Cohen & D’Esposito, 2016; Genon et al., 2018) at different topological levels (i.e., whole-brain/system-level, modular/subsystem-level, and region/nodal-level) (Farahani et al., 2019; Fornito et al., 2016). Across the lifespan, cognitive efficiency is supported by a balance between integrated and modular information processing (Bullmore & Sporns, 2012; Meunier et al., 2010; van den Heuvel & Sporns, 2013). In other words, optimal coordination of neural activity is based on global homeostasis – the ability to adapt and maintain stability in the face of changing conditions (see also the notion of *metastability*: Naik et al., 2017; Tognoli & Kelso, 2014).

In line with our previous findings (Roger, Rodrigues De Almeida, et al., 2022), a recent systematic review of resting-state data studies reported that reduced local efficiency at the system level, along with reduced segregation and enhanced integration within and between RSNs at the subsystem level, are the connectomic fingerprints of healthy aging with an inflection point in midlife (Deery et al., 2023). However, a crucial challenge resides in understanding the neural mechanisms that bridge reduced segregation with enhanced integration (Stumme et al., 2020), and how these mechanisms induce a neurocognitive dynamic that reflects the changes in cognitive performance as age advances.

Indeed, studies report contradictory findings depending on the topological level of analysis. At a system level, age-related enhanced integration (Battaglia et al., 2020) would be generally associated with a dedifferentiated system that fails to alternate efficiently between integrative and segregation states of connectivity (Chan et al., 2014; Chan et al., 2017), thus highlighting a maladaptive process. At a subsystem or brain network level, brain regions would undergo similar dedifferentiation processes translating to reduced functional specialization (Goh et al., 2010; Park et al., 2004). This is primarily reflected by reduced segregation and enhanced integration in the sensorimotor and higher-order associative and control networks (Deery et al., 2023; Wig, 2017). However, in contrast with the system level, enhanced integration has been shown to be compensatory (Cabeza et al., 2018) with direct benefits for cognitive efficiency (Bertolero et al., 2015; Meunier et al., 2010). Specifically, increased coupling between default-mode (DMN) and fronto-parietal (FPN) networks correlated with better global cognitive performance (Spreng et al., 2018; Spreng & Turner, 2019). Moreover, it appears that the ability to deactivate DMN regions may be a key ingredient of cognitive resilience (capacity to mitigate the deleterious effect of dedifferentiation) across the lifespan (Deery et al., 2023; Grady et al., 2016; Singh-Manoux et al., 2012; Varangis et al., 2019). Indeed, older adults show reduced DMN deactivation (Spreng & Turner, 2019), potentially impacting the interaction between semantic and control-executive processes, as observed for LP (Baciu et al., 2021) and verbal fluency (Muller et al., 2016; Whiteside et al., 2016).

The main objective of this study is to investigate how the age-related reorganization of the language connectome explains the discrepancy between LC and LP performances across the lifespan. By assuming that language is a complex function working in synergy with long-term memory and executive processes, we aimed to model the neural mechanisms that support this inter-cognitive functioning. To address this objective, we leveraged a population-based resting-state fMRI dataset from the CamCAN cohort (Cam-CAN et al., 2014) and applied graph theory analyses to evaluate the reorganization of the language connectome in terms of integration and segregation properties at multiple topological scales of analysis.

## 2. Materials & Methods

### 2.1. Participants

We included 652 healthy adults from the Cambridge Center for Ageing and Neuroscience project cohort (Cam-CAN et al., 2014). Further recruitment information can be found in Taylor et al. (2017). Our analysis focused exclusively on functional fMRI brain data obtained during a resting state period. After careful examination, we excluded 24 participants and had a final sample size of 628 participants (age range: 18-88; 320 females; 208 males). The 24 participants were excluded from the analysis for various reasons, including missing functional imaging data (N = 4), incomplete volume acquisition (N = 4; however, one of the four subjects has been retained with 74% (194/261) of the total number of volumes), unreliable CSF mask (N = 9), and having 10% or more outlying volumes after preprocessing (M = 13.6, SD = 3.2, N = 8). We chose eight cognitive tasks for brain-cognition analyses to span a continuous spectrum from a high to a low degree of synergy between language, long-term memory, and executive functions (Table 1). The sample of 628 participants was further reduced to 613 due to missing cognitive data. Specifically, we excluded participants with more than 3 missing neuropsychological scores (N = 15). Then, we replaced any missing score of the remaining 613 participants with the appropriate median of their age decile.

**Table 1.** General presentation of the eight cognitive tasks. A detailed description of each task is presented in Table S1 and Appendix S1.

| **Cognitive Task** | **Processes involved**<br>(from high to low amount of inter-cognitive synergy) |
| --- | --- |
| Naming<br>(Clarke et al., 2013) | Language (phonological access & semantic memory) |
| Verbal Fluency<br>(Lezak et al., 2012) | Language (phonological, semantic) & Executive function (EF) |
| Proverb comprehension<br>(Huppert et al., 1994) | EF (abstraction) & Language (comprehension) |
| Tip-of-the-tongue<br>(R. Brown & McNeill, 1966) | Language (retrieval) & EF (error monitoring) |
| Hotel task<br>(Shallice & Burgess, 1991) | EF (planning & multitasking) |
| Cattell task<br>(Cattell & Cattell, 1960) | EF (Fluid intelligence) |
| Story recall<br>(Tulsky et al., 2003) | Long-term memory |
| Sentence comprehension<br>(Rodd et al., 2010) | Language (syntactic, semantic) |

### 2.2. MR acquisition

For information regarding the MR acquisition and the resting state protocol applied in this study, please refer to Appendix S1. Further details are provided by Cam-CAN et al. (2014), as the data was sourced from the Cambridge Center for Ageing and Neuroscience project cohort.

### 2.3. Resting-state fMRI data analysis

#### 2.3.1. Data Preprocessing

The rs-fMRI data underwent preprocessing using SPM12 (Welcome Department of Imaging Neuroscience, UK, http://www.fil.ion.ucl.ac.uk/spm/) within MATLAB R2020b (MathWorks Inc., Sherborn, MA, USA). We employed a standard preprocessing pipeline (including realignment, reslicing, co-registration, segmentation, normalization, and smoothing) similar to that described in our previous work (see Roger et al., 2020) with specific details mentioned in Appendix S1.

#### 2.3.2. Cerebral parcellation: the LANG connectome atlas

The LANG connectome atlas, referred to as the LANG atlas, comprises a collection of 131 regions of interest (ROI) derived from a panel of 13 language fMRI tasks (see Roger, Rodrigues De Almeida, et al., 2022). Each ROI is represented by a spherical region with a diameter of 6 mm, centered on the MNI coordinates proposed by Power et al. (2011). To transform the LANG atlas, which is based on extrinsic fMRI activation, into a resting-state LANG (rs-LANG) atlas consisting of ROIs derived from the intrinsic activity, we labeled all LANG regions according to their primary resting-state network (RSN) based on the Cole-Anticevic Brainwide Net Partition (CAB-NP; Ji et al., 2019). We utilized a publicly available volumetric version of the CAB-NP that was converted to standard MNI space. By overlaying the volumetric RSN map onto the 131 regions (see Banjac et al., 2021), we determined the number of voxels overlapping each region and each RSN. This approach ensured accurate voxel-based labeling of each region to their primary RSN (see Appendix S3 for detailed results). When examining the modular organization of rs-LANG (see Figure 1), the RSN composition of a given subsystem was weighted by the mean percentage of overlap between each region and their primary RSN. This improvement led to a 12% increase in mean accuracy.

**Figure 1.**
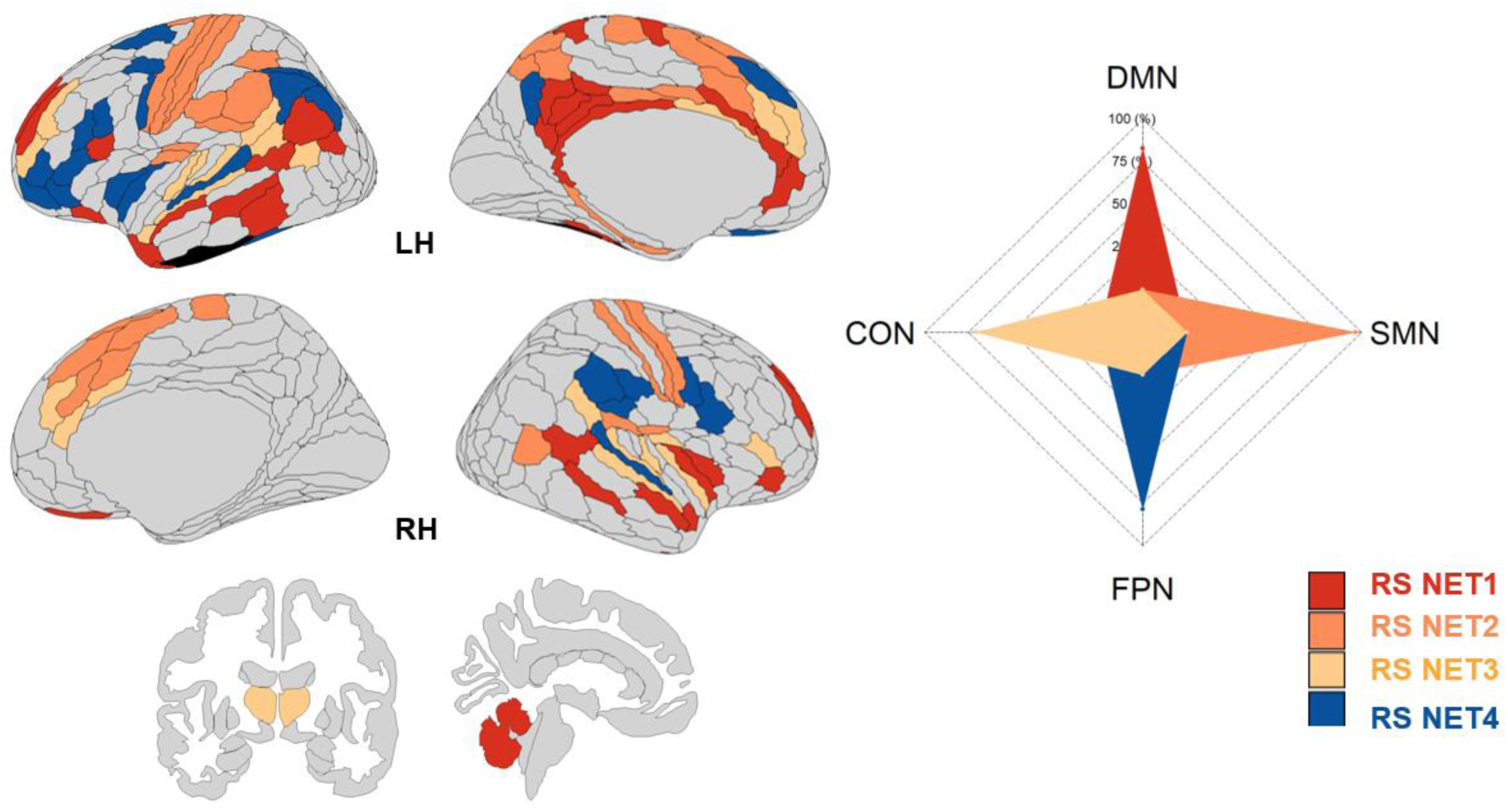
Illustration of the modular organization of the language connectome at rest. Each RS NET is a subsystem obtained from the consensus clustering procedure (see the method section 2.4.2). An additional 3-region module (black) associated with the VMM (Ventral Multi-Modal) network was also identified but not considered for analysis due to its instability as a stand-alone module across the lifespan. *Abbreviations: LH (left hemisphere); RH (right hemisphere);* RS NET1 (40 regions, red), RS NET2 (34 regions, orange), RS NET3 (32 regions, yellow), RS NET4 (22 regions, blue). DMN (Default Mode Network); FPN (Fronto-Parietal Network); CON (Cingulo-Opercular Network); SMN (Sensorimotor Network). Brain visualization was done with the package *ggseg* in R (Mowinckel & Vidal-Piñeiro, 2020) and projected on a multimodal cortical (HCP_MMP1.0; Glasser et al., 2016) and subcortical parcellation (Fischl et al., 2002). For details, please refer to Figure S1 and S2 in Appendix S2, and Appendix S3.

#### 2.3.3. Resting-state LANG connectomes

Using the CONN toolbox (version 21.a; Nieto-Castanon, 2020), we conducted an ROI-to-ROI analysis and generated a connectivity matrix of dimensions 131x131 for each participant using Fisher-transformed bivariate correlation coefficients. These connectivity matrices were subsequently employed for network analysis. We disregarded negative correlations by setting them to zero, consistent with previous studies (Chong et al., 2019; Martin et al., 2022; Wang et al., 2011). Additionally, we applied thresholding to each matrix at five sparsity levels (10, 12, 15, 17, and 20%). This step aimed to reduce the presence of spurious connections and was based on the most likely sparsity levels known to produce a small-world organization (see Appendix S2), as outlined by Achard and Bullmore (2007). Correspondingly, the thresholded matrices were binarized to generate five undirected graphs for each participant.

### 2.4. Resting-state LANG network analysis

The network or graph analysis measured the information flow within each connectome (Rubinov & Sporns, 2010). Specifically, we assessed (i) the balance between functional integration and segregation at the system level, (ii) the modularity at the subsystem level, and (iii) the information transfer at the nodal level by evaluating the topological role of each region.

#### 2.4.1. System-level analysis: integration vs. segregation balance assessment

Using the Brain Connectivity Toolbox (BCT) implemented in MATLAB 2020b and available at https://www.nitrc.org/projects/bct/ (Bullmore & Sporns, 2009), we extracted three key graph metrics: (1) global efficiency (E_glob_), (2) local efficiency (E_loc_), and (3) clustering coefficient (Clust_coeff_). Globally, E_glob_ was calculated as the inverse of the shortest path lengths or the average of unweighted efficiencies across all pairs of nodes (Latora & Marchiori, 2001). This metric quantifies the efficiency of parallel information transmission across the global network (Bullmore & Bassett, 2011). Locally, E_loc_ is similar to E_glob_ but on node neighborhoods. When averaged at the system level, it illustrates the segregation property of a network in processing information (Latora & Marchiori, 2001). For each node, the clustering coefficient (Clust_coeff_) was calculated as the fraction of a node’s neighbors that are also neighbors of each other (Rubinov & Sporns, 2010), and averaged at the system level. We visually inspected E_glob_ and Clust_coeff_ across the five sparsity levels and determined that the optimal threshold that balances integrated and modular processing was 15% (Figure S3, Appendix S2). We further ensured that this threshold was optimal by repeating the procedure for each age decile. Following this, we reduced all graphs to a fixed number of edges by retaining the top 15% (2554 edges). Additionally, we verified that the reduced connectomes maintained full connectivity and were sufficiently devoid of isolated nodes, meaning that the largest connected component (LCC) included at least 80% of all nodes. After thresholding at 15%, we characterized the balance between functional integration and segregation at the system level by determining the relative predominance of global (E_glob_) versus local efficiency (E_loc_). A higher balance reflects a higher predominance of integration or higher integrative/global efficiency.

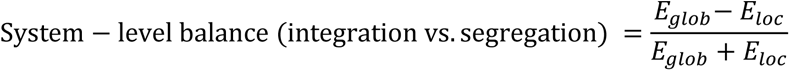

#### 2.4.2. Subsystem-level analysis: modularity assessment

To examine the modular organization of rs-LANG, we employed the Louvain community detection algorithm (Blondel et al., 2008) with a resolution parameter set to γ = 1.295, aiming to align with the RSNs identified in the CAB-NP Atlas (Ji et al., 2019). As different runs of the algorithm can yield varying optimal partitions, we implemented a consensus clustering approach (Lancichinetti & Fortunato, 2012). This approach involves iteratively clustering co-assignment matrices until convergence, aggregating the frequency of node assignments to the same module. To reduce spurious node assignments, we applied a threshold of τ = 0.5 to the co-assignment matrices (Jiang et al., 2021). If a node did not consistently belong to the same module in at least 50% of the iterations, its co-assignment weight was set to zero. We executed the algorithm 1000 times for each subject and repeated the entire process 1000 times to generate a group-level consensus partition based on the subject-level partitions. We ensured that this consensus partition accurately represented all individuals across different ages, confirming its suitability for statistical analysis (refer to Appendix S2 for further information).

#### 2.4.3. Nodal-level analysis: topological roles

At the nodal level, we investigated the topological reorganization of each connectome by defining four distinct topological roles: Connector, Provincial, Satellite, and Peripheral (Bertolero et al., 2015; Guimerà & Amaral, 2005). Therefore, the composition of each individual’s language connectome is represented by four percentages which add up to 100%. To assign each role to a node (i.e., a region), we employed two graph metrics: (i) the Within-Module Z-score (WMZ; Latora & Marchiori, 2001), which quantifies connectivity within subsystems (short-range), and (ii) the normalized Participation Coefficient (PC; Pedersen et al., 2019) which measures connectivity between subsystems (long-range) while removing the bias associated with the number of nodes in each module. Consistent with prior research (Roger, Rodrigues De Almeida, et al., 2022; Schedlbauer & Ekstrom, 2019), we standardized both metrics (WMZ and zPCnorm) for each individual and assigned a topological role to each node. In relation to the entire set of 131 regions, a Connector node displays a high proportion of both short- and long-range connections (zPCnorm >= 0, WMZ >= 1e-5). A Provincial node displays a high proportion of short-range connections (zPCnorm < 0, WMD >= 1e-5). A Satellite node displays a high proportion of long-range connections (zPCnorm >= 0, WMZ < 1e-5). A Peripheral node, on the other hand, is functionally withdrawn from the network (zPCnorm < 0, WMZ < 1e-5).

### 2.5. Statistics

We conducted statistical analysis in two steps. First, at a connectomic level, we examined the evolution of the relative proportion of topological roles across the lifespan (i.e., quantitative analysis). Subsequently, we modeled the neural mechanisms driving this evolution using a probabilistic framework (i.e., qualitative analysis). This approach allowed us to uncover the patterns and principles governing the age-related connectivity changes in the language connectome. Second, at a neurocognitive level, we employed canonical correlation analysis (CCA) to examine the many-to-many relationships between these neural mechanisms and the changes in cognitive performances across the lifespan. We controlled for mean FC, gender, and total intracranial volume in all models (Eikenes et al., 2023). Mean FC was calculated by averaging the positive weights of the unthresholded upper triangular connectivity matrix.

#### 2.5.1. Connectomic dynamic across the lifespan

To evaluate the age-related topological changes of the language connectome, we examined how the relative proportions of four topological roles (connector, provincial, satellite, peripheral) evolve across the lifespan. Due to the inherent limitations of percentage-based statistics (which sum up to 100%), namely, high correlation and dependence on pairwise covariance – we applied a log-based transformation to the data (Smith et al., 2016). This transformation, analogous to a log odds transformation, removes the lower (0%) and upper (100%) boundaries of the original metric (i.e., removing the unit-sum constraint), thus remaining relatively easy to interpret:

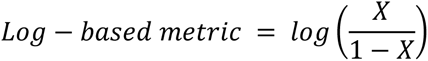

Here, X represents the percentage-based proportion of either Connector, Provincial, Satellite, or Peripheral nodes. To handle undefined logarithms for zero entries, we imputed percentages using Bayesian non-parametric multiplicative replacement with the package zCompositions in R (Martín-Fernández, 2003; Palarea-Albaladejo & Martín-Fernández, 2015).

##### Quantitative analysis

Following this transformation, the effect of age was examined using generalized additive models (GAM; *mgcv* package in R; Wood, 2006, 2017) at a system and subsystem level, using a 3-knot spline to mitigate overfitting concerns. Factor-smooth interactions were applied at a subsystem level, and p values were corrected at the False Discovery Rate (*q < 0.05*).

##### Qualitative analysis

To elucidate the neural mechanisms driving quantitative topological changes in the language connectome, we identified the most likely topological role that a region may occupy within younger (18-44), middle-aged (45-55), and older (>55) individuals. We chose these age groups to stay consistent with the quantitative results reported in section 3.2.

This involved 3 steps. (i) First, we determined the frequency at which each region is assigned each topological role for each age group. For example, among younger adults, a region may be considered a connector node for 75% of participants but a satellite node for the remaining 25%. (ii) Second, we took the outer product of all the younger and middle-aged frequency values to calculate the probability of all 16 possible trajectories between roles. For example, if a region has a 75% probability of being a provincial node in younger adults and 30% a connector node in middle-aged adults, then the resulting provincial-to-connector trajectory has an occurrence probability of 22.5% (0.75*0.3). (iii) Third, we repeated this calculation between these 16 trajectories and the frequency values and the older age group (16*4 = 64 trajectories in total).

Finally, we selected the most likely trajectory of a region as the one with the highest probability (e.g., provincial-to-connector-to-connector). To account for the inter-individual variability within each age group, we also included the trajectories whose probability fell within a 5% range from the highest one for each region. Using this approach, we captured the most likely transitions of topological roles between age groups for each region while also considering the variability within each age group.

#### 2.5.2. Neurocognitive Dynamic: Canonical Correlation Analysis

To identify the many-to-many relationships between brain functional connectivity changes and neuropsychological scores, we conducted a canonical correlation analysis (CCA). CCA works by finding the linear combinations within the brain and cognitive set of variables that maximize the correlation between the two sets (Wang et al., 2020).

##### CCA setup

We organized the data into a brain-functional (X) and cognitive (Y) matrix to set up the CCA. For X and Y, scores were z-scored. Additionally, we prepended two orthogonal contrasts in the cognitive dataset that we set between three age groups: (contrast #1) 56-60 > 45-50 + 51-55, (contrast #2) 51-55 > 45-50. These contrasts ensured that the model could also account for nonlinear relationships with age reported in section 3.2. CCA analysis produces as many canonical functions as the number of variables within the smaller set (i.e., 8 cognitive scores + 2 age contrasts). Each canonical function is composed of a cognitive and a brain variate comparable to latent variables (Sherry & Henson, 2005). The robustness of the results was assessed using a 10-fold cross-validation with 1,000 bootstrap resamples.

##### CCA interpretation

At a cognitive level, we report CCA results using structure coefficients (*r*) – the correlation between an observed variable and its corresponding variate. Thus, the higher the correlation, the greater the contribution of said variable to said variate. At a neurocognitive level, given that the neural mechanisms represent our unit of interest, we computed the difference between the structure coefficients associated with each variable of a given mechanism. Considering a reconfiguration from a provincial to a connector role as an example, we subtracted the provincial variable’s structure coefficient from the connector one. Importantly, we reported the cross-correlations between the brain variables and the cognitive variates. The resulting coefficient served as a proxy for the correlation between said mechanism and said cognitive variate, providing an intuitive understanding of how a neural mechanism affects age-related cognitive performance.

## 3. Results

### 3.1. Modular organization of the Language connectome at rest (rs-LANG)

Applying modularity analysis to the resting-state Language connectome (rs-LANG) uncovered four main subsystems. Figure 1 provides a visual representation of these subsystems. Appendix S2 provides a detailed comparison with the task-based modular organization of the same connectome, described by Roger, Rodrigues De Almeida, et al. (2022).

Considering the composition, the largest subsystem, RS NET1 (40 regions), comprises 66.4% of DMN regions involved in higher-level cognitive function and can thus be regarded as the associative subsystem. The second largest subsystem, RS NET2 (34 regions), is saturated by sensorimotor regions (SMN; 77.1%) along with contributions from CON regions (15.3%) and can thus be regarded as the sensorimotor subsystem. The third largest subsystem, RS NET3 (32 regions), engages the cingulo-opercular network (CON; 56.4%), which can thus be regarded as the bottom-up attentional subsystem (Dosenbach et al., 2024; Wallis et al., 2015). The smallest subsystem, RS NET4 (22 regions), is saturated by FPN regions (63.1%), with nontrivial contributions from CON regions as well (17.2%), and can thus be regarded as the top-down control-executive subsystem. Interestingly, all 11 regions of the conventional language network defined by Ji et al. (2019) coalesce with the associative (5 regions) and bottom-up attentional subsystems (4 regions) while the remaining 2 regions form crucial short-range connections with the top-down control-executive subsystem. This suggests that core language processing at rest is functionally clustered with subsystems that support both semantic access and top-down/bottom-up cognitive control (see Appendix S3 for details).

### 3.2. Connectomic dynamics of the language connectome

#### Quantitative changes

At the system level, we found that healthy aging is associated with reduced local efficiency (*t* = -10.08, *p* < .001, *η*_2p_ = .15, 95% CI [.11; Inf]) and preserved global efficiency (*p* = .53), which tilts the balance between integration and segregation towards a higher integrative efficiency as age increases (*t* = 9.4, p < .001, *η*_2p_ = .13, 95% CI [.09; Inf]) (Figure 2A). We also found that healthy aging is associated with a decrease in the proportion of provincial nodes (*b* = -0.08 95% CI [-0.1; -0.05]; F(1, 624) = 47.2; *p* < .001; *η*_p_^2^ = .07) and satellite nodes (*b* = -0.02 95% CI [-0.03; -0.01]; F(1, 624) = 4.7; *p* = .03; *η*_p_^2^ = .01), which is contrasted by an increase in the proportion of connector nodes (*b* = 0.05 95% CI [0.03; 0.06]; F(1, 624) = 27.9; *p* < .001; *η*_p_^2^ = .04) and peripheral nodes (*b* = 0.03 95% CI [0.02; 0.05]; F(1, 624) = 16.8; *p* < .001; *η*_p_^2^ = .03) (Figure 2B).

**Figure 2.**
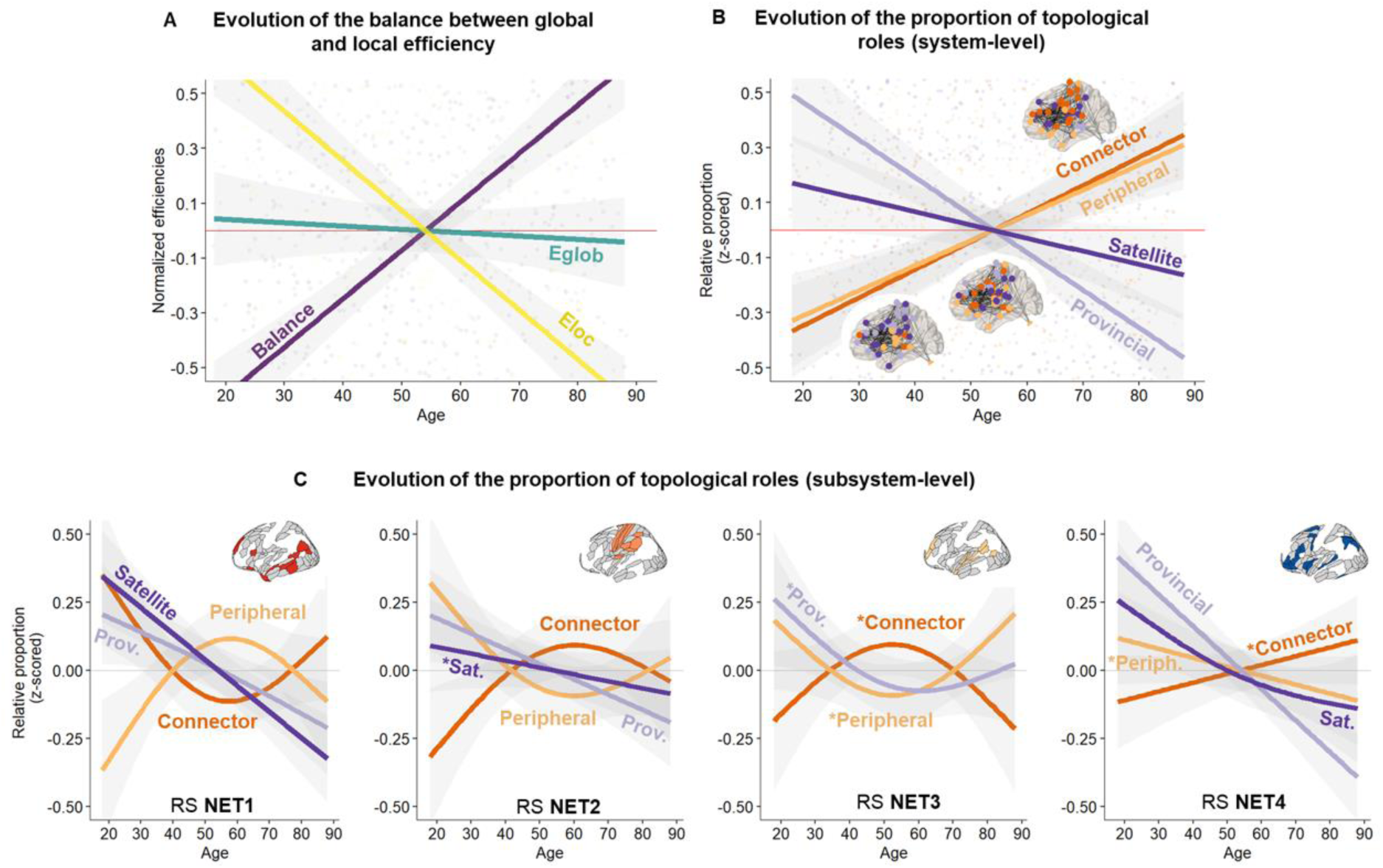
System-level topological dynamics across the lifespan. **(A)** Illustrates normalized efficiencies (y-axis) as a function of age (x-axis). Eloc = Local efficiency; Eglob = Global efficiency; Balance = dominance of global efficiency calculated as (Eglob – Eloc) / (Eglob + Eloc). **(B and C)** Evolution of the relative proportion of topological roles (y-axis) with age (x-axis). For the subsystem level (Panel C), changes with a tendential statistical significance are indicated by a star next to the label. *Abbreviations:* RS NET (subsystems of the language connectome at rest); RS NET1 (Associative); RS NET2 (Sensorimotor); RS NET3 (Bottom-up attentional); RS NET4 (Top-down control-executive). Please refer to Figure 1 for the composition of each RS NET. Connector (High integration/High segregation); Provincial (Low integration/High segregation); Satellite (High integration/Low Segregation); Peripheral (Low integration/Low segregation).

At the subsystem level (Figure 2C), we found two patterns of coordinated changes between subsystems: (i) First, we observed that a major loss of satellite nodes in RS NET1 was coordinated with a major loss of provincial nodes in RS NET4 (F = 22.05/28, *p* < .001/.001, *edf* = 1/1), and conversely with more moderate losses (F = 6.07/3, *p* = .015/.042, *edf* = 1.29/1.32). Of note, we also found a moderate loss of provincial nodes in RS NET2 (F = 6.8, *p* = .009, *edf* = 1). (ii) Second, we observed coordinated nonlinear changes in the proportion of connector (u-shape) and peripheral nodes (inverted u-shape) in RS NET1 with an inflection point at age 55 (F = 3.61/3.8, *p* = .02/.02, *edf* = 1.88/1.87). Interestingly, we found the same anticorrelation pattern in RS NET2 (F = 2.7/2.8, *p* = .047/.045, *edf* = 1.78/1.79) but this pattern was mirrored compared to the one observed in RS NET1 and highlighted an inflection point at age 60.

Of note, several tendential associations with age suggest that some mechanisms could be underpinned by (unmodeled) age-invariant factors: a linear decrease of satellite nodes in RS NET2 (*p* = .23); a coordinated pattern between connector (*p* = .08) and provincial/peripheral nodes in RS NET3 (*p* = .13/.09); a linear increase and decrease respectively in connector (*p* = .12) and peripheral nodes (*p* = .18) in RS NET4.

In sum, analyses at the system and subsystem level revealed that midlife is a critical period for functional brain reorganization of the language connectome. Specifically, changes in RS NET1 are pivotal: (i) the loss of satellite nodes in RS NET1 is coupled with the loss of provincial nodes in RS NET4; (ii) the amount of connector and peripheral nodes are anticorrelated within a given subsystem, and this anticorrelation pattern tends to be mirrored between RS NET1 and the other subsystems (see Figure 2C).

#### Qualitative changes

To clarify the mechanisms driving the topological changes reported above, we conducted a probabilistic analysis. Overall, we found that 35.1% (46 regions) of the language connectome undergoes a topological reorganization, suggesting that some regions occupy different roles throughout life. Across subsystems, 50% of RS NET4, 50% of RS NET2, and 34% of RS NET1 regions reconfigure, whereas regions in RS NET3 are more inflexible (24.3%). A web app is available to explore the reconfiguration cross-sectionally: https://lpnc-lang.shinyapps.io/seneca/. To account for inter-individual variability, we consider 17 additional trajectories (see section 2.5.1 for details on the calculations), which amounts to 63 (46 regions that reconfigure + 17) trajectories.

In line with the previous results, healthy aging was associated with a substantial gain of connector (53.9% of all reconfigurations; 34 regions) and peripheral nodes (19%; 12 regions). We observed that this topological reorganization was achieved in two ways: either (i) reallocating short-range connections within subsystems or (ii) reallocating long-range connections between subsystems. Figure 3 depicts these two dynamics. Table S1 and S2 in Appendix S2 summarize the following results.

**Figure 3.**
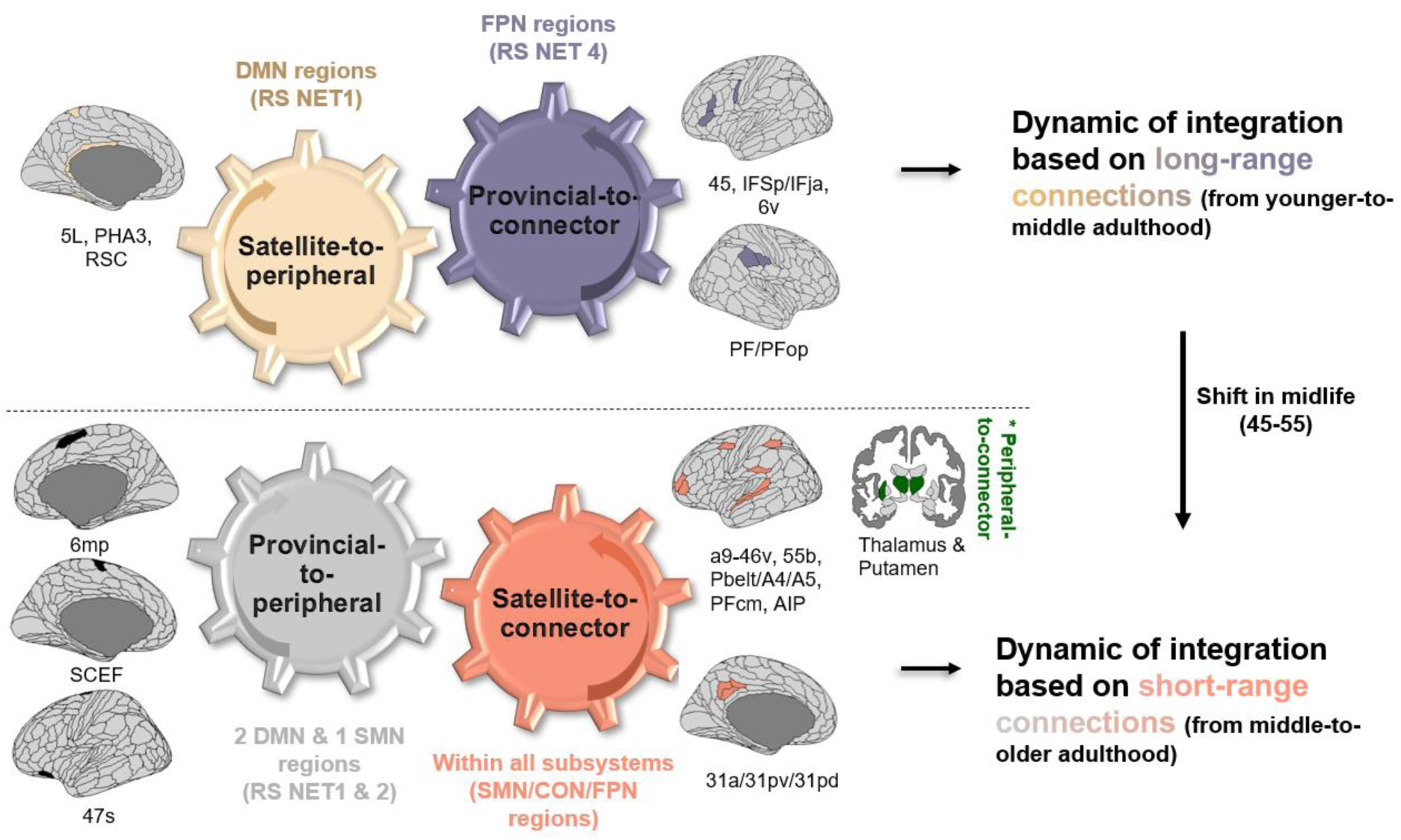
Probabilistic topological model of age-related integration. Two dynamics of integration across the lifespan are proposed: (i) “energy-costly” dynamic based on long-range connections between subsystems, highlighting the flexible DMN-FPN coupling in younger adulthood, and (ii) “energy-efficient” dynamic based on short-range connections within subsystems, highlighting a less flexible DMN-FPN coupling in older adulthood. *Abbreviations:* RS NET (subsystems of the language connectome at rest); RS NET1 (Associative); RS NET2 (Sensorimotor); RS NET3 (Bottom-up attentional); RS NET4 (Top-down control-executive). Please refer to Figure 1 for the composition of each RS NET. Connector (High integration/High segregation); Provincial (Low integration/High segregation); Satellite (High integration/Low Segregation); Peripheral (Low integration/Low segregation). DMN (Default Mode Network); FPN (Fronto-Parietal Network); CON (Cingulo-Opercular Network); SMN (Sensorimotor Network). Labels under brain illustrations are the names of the regions following the labeling proposed by Glasser et al. (2016). Brain visualization was done with the package *ggseg* in R (Mowinckel & Vidal-Piñeiro, 2020) and projected on a multimodal cortical (HCP_MMP1.0; Glasser et al., 2016) and subcortical parcellation (Fischl et al., 2002).

On the one hand, we observed a dynamic governed by the loss and gain of new short-range connections via provincial-to-peripheral (9.5% of all trajectories; 6 regions) and satellite-to-connector reconfigurations (20.6%: 13 regions). From early to middle adulthood, we observed that RS NET1 (right MTG; 92% FPN, right BA44; 72% DMN) and RS NET4 (left MFG; 99% FPN) are the most likely to lose these connections, while from middle to older adulthood, this impacted RS NET2 (right SMA; SMN, left paracentral lobule; 74% DMN) and RS NET1 (left insula; 72% DMN). Interestingly, we note that the integration of these short-range connections occurs mainly from middle to older adulthood (69.2%; 9 out 13 regions) within all subsystems: in the left middle cingulate cortex (RS NET1; 11.1%), left supramarginal gyrus and left postcentral area (RS NET2; 22.2%), left superior temporal gyrus (RS NET3; 11.1%), and left precentral/middle frontal regions in RS NET4 (22.2%). Additionally, we noticed the key role of subcortical areas (33.3%; left/right thalamus, left putamen; RS NET3) which also undergo unique peripheral-to-connector reconfigurations (left and right putamen) from middle to older adulthood. Given that changes in peripheral/connector proportions in RS NET3 were only tendentially related to age in the previous section, this suggests that integration via these subcortical regions could also be dependent on age-invariant factors.

On the other hand, we found a dynamic governed by the loss and gain of long-range connections via satellite-to-peripheral (9.5% of trajectories; 6 regions) and provincial-to-connector reconfigurations (30.1%: 19). While the integration is driven by FPN regions in RS NET 4 (45%) and in left frontal/precentral/postcentral regions (SMN) in RS NET2 (55%) from early to middle adulthood, we noticed that it is almost exclusively implemented by RS NET2 from middle to older adulthood (SMN: SMA and pre/post-central regions). Interestingly, this coincides with our previous observation that some FPN regions in RS NET4, which integrate long-range connections in early adulthood, shift to short-range connections in older adulthood (i.e., left MFG and left precentral). Interestingly, this shift co-occurs with a similar transition in RS NET 1 in middle age (age 45-55). Indeed, while RS NET1 undergoes a “deactivation” process (i.e., satellite-to-peripheral) from early to middle adulthood, losing long-range connections (left fusiform area DAN/DMN; left paracentral lobule DMN; left pCC DMN; 83.3% of all deactivations), it begins integrating connections from middle to older adulthood.

Our results also show that short-range connections may become increasingly more scarce as age increases, compromising both dynamics of integration. Indeed, we observed that (i) regions in RS NET1 (left IFG DMN; left aCC CON/DMN), RS NET2 (left SFG CON/SMN; right Rolandic Op SMN/CON), and RS NET 4 (left insula FPN/CON) are likely to lose the short-range connections integrated earlier in life (i.e., connector-to-satellite reconfigurations), but also that (ii) the number of provincial-to-satellite reconfiguration increases, showing that the older adult brain is less likely to maintain connector properties especially in right superior frontal areas in RS NET1 (DMN) and left parahippocampal lobule in RS NET2 (SMN).

In sum, our observations suggest that the shift reported in midlife (45-55) is triggered by a reduced deactivation in RS NET1 (i.e., satellite-to-peripheral) coupled with a reduced integration of long-range connections in RS NET4 (i.e., provincial-to-connector).

### 3.3. Neurocognitive Dynamics: Canonical Correlation Analysis

To evaluate the neural mechanisms that support healthy cognitive aging, we employed canonical correlation analysis (CCA). CCA yielded two significant canonical functions, each composed of a cognitive and brain variate that maximize the correlation between the brain and cognitive set of variables (Wilks’ Λ = .68, *R_c_*_1_ = .33; *R_c_*_1_^2^ = 11%, *p* < .001; Wilks’ Λ = .76, *R_c_*_2_ = .27; *R_c_*_2_^2^ = 7%, *p* = .04). Below, we first describe the results at a cognitive level and then at a neurocognitive level.

#### Cognitive level

Figures 4A & 4C show that Variate I was associated with a steady decline with age (F = 388.7, *p* < .001, *edf* = 1.51; *r_constrast_older > middle + younger_* = -.71, *r_constrast_middle > younger_* = -.49). This primarily impacted performances in language-related tasks recruiting executive functions such as multitasking (*r* = .56), lexical production (.49), fluid intelligence (i.e., Cattell = .48), verbal fluency (.48), tip-of-the-tongue (.38) and semantic abstraction (i.e., proverb task = .34) (see Figure 4A and 4B). Variate II was nonlinearly associated with age (F = 90.1, *p* < .001, *edf* = 1.99; *constrast_middle > younger_* = .62), marked by a transition in midlife in line with our previous findings. Interestingly, this correlated with better overall semantic performances (semantic abstraction = .43; sentence comprehension = .21), which was also associated with better naming (.40) and marginally better verbal fluency (.11). These changes were also proportional to increased fluid-related challenges (-.32), suggesting that heightened semantic knowledge in midlife could help maintain lexical production abilities in the face of fluid processing decline.

**Figure 4.**
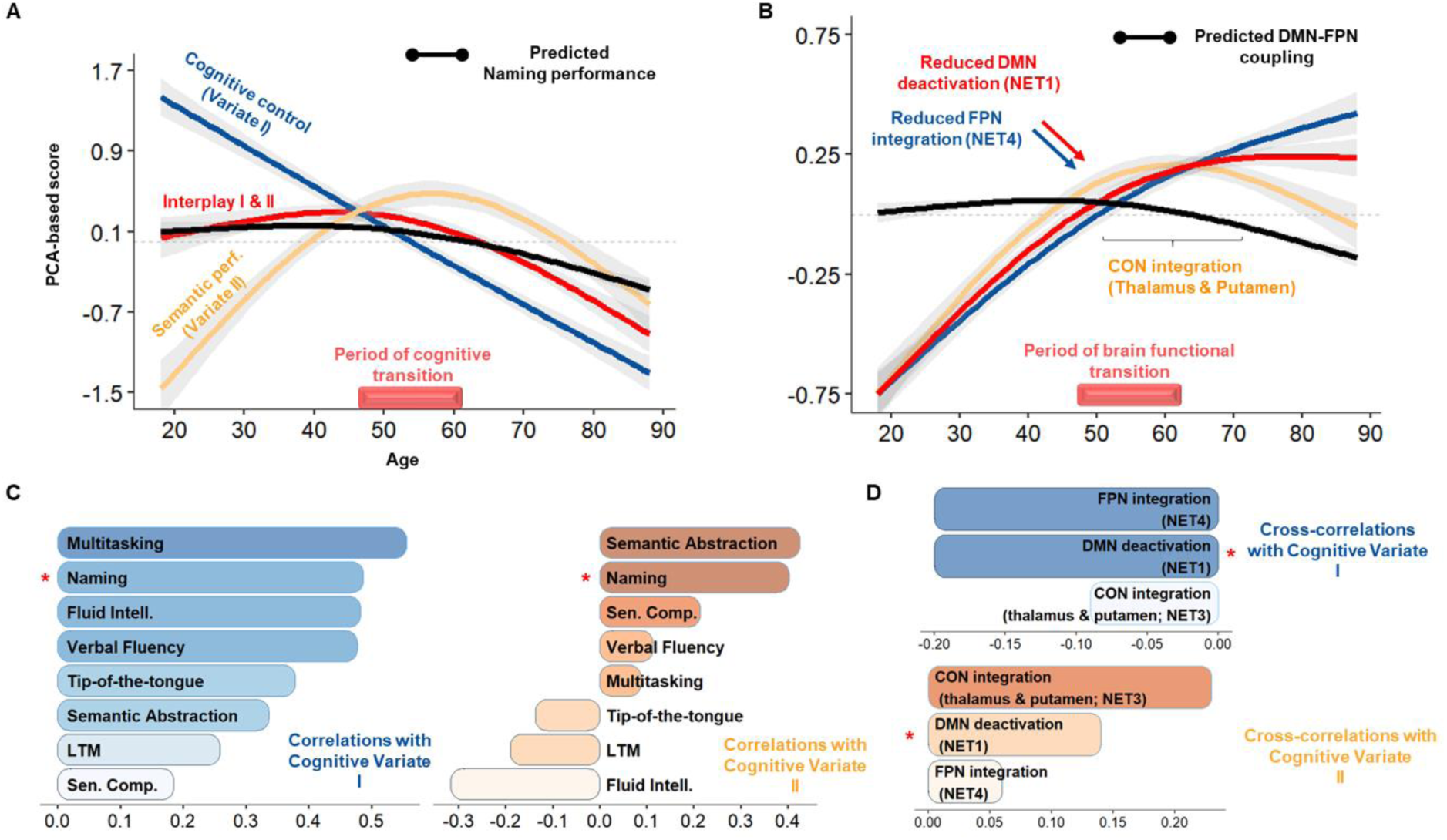
Age-related neurocognitive dynamics of language. **(A and B)** Illustrate the age-related trajectory of the cognitive variate and the corresponding neural mechanisms according to the results. The bar above the x-axis reports age points with a significant second-order derivative for most trajectories, reflecting a neurocognitive transition in midlife. **(C and D)** Illustrate the structure coefficients with the cognitive variates, the correlations (cognitive variable-cognitive variate), or cross-correlations (brain mechanism-cognitive variate). The red star indicates that naming and DMN suppression are highly covariant with each variate, thus showing that DMN suppression underpins naming performances during healthy aging.

Lexical production (naming) was highly correlated with both variates (see Figure 4C). Verbal fluency and semantic abstraction were also highly correlated with both variates, although the former was preferentially correlated with cognitive control (Variate I) and the latter with semantic performance (Variate II). This suggests that age-related decline in these tasks (see Table 1, section 2.1) stems from cognitive control and semantic cognition (i.e., semantic control).

#### Neurocognitive level

Functional deactivation (i.e., gain in peripheral nodes) and functional integration (i.e., gain in connector nodes) were largely anticorrelated with Variate I. This suggests that the dynamics of integration based on the reallocation of long or short-range connections may help mitigate executive function deficits as age increases.

We also found that the joint trajectory of DMN deactivation and FPN integration fit the predicted trajectory of naming performances (t(611) = 245.15, *p* < .001):

i. The loss of long-range connections of RS NET1 regions (i.e., DMN deactivation) was both anticorrelated with Variate I (-.20) and correlated with Variate II (.12), indicating that this mechanism is crucial for mitigating fluid processing decline with heightened semantic representations as age increases. Considering the cognitive changes reported above, this certainly contributes to delaying the onset of difficulties in naming, verbal fluency, and semantic abstraction. In line with this, from middle to older adulthood (see Figure 4B & 4D), high correlation with Variate II (i.e., above the median) confirms that reduced deactivation (0.14), and correspondingly enhanced functional integration of long-range connections in RS NET1 (−0.09), compromises the ability to compensate fluid processing decline (-.32) with semantic knowledge (i.e., reduced semantic abstraction; proverb task = .43), with implications on naming performances (.40).
ii. The integration of long-range connections in RS NET4 (i.e., FPN regions) had a comparable effect. Nonetheless, it was preferentially correlated with executive function mitigation (Variate I: -.22; Variate II: .06). Considering the cognitive changes reported above, this mechanism may primarily enhance verbal fluency and the cognitive control component of lexical production (naming).

Additionally, increased peripheral-to-connector reconfigurations in RS NET3 (0.23) were mostly associated with Variate II (.23). In line with our probabilistic results, this mechanism peaked slightly after midlife (see Figure 4B). This suggests that, despite reduced DMN deactivation, integration in the bottom-up attentional system (RS NET3), specifically in the bilateral thalamus and left putamen (please refer to section 3.2), could promote semantic abstraction in addition to the semantic component of lexical production (naming).

## 4. Discussion

Healthy aging is associated with a heterogeneous decline across cognitive functions, including language. Specifically, language production (LP) declines more rapidly than language comprehension (LC). The neural mechanisms underlying this variability still need to be understood. In this study, we leveraged resting-state fMRI and neuropsychological data from the population-based CamCAN cohort (Cam-CAN et al., 2014) to investigate the functional reorganization of the language connectome and its association with age-related cognitive variability. Employing state-of-the-art graph theoretical analysis, we developed a data-driven pipeline that integrates both cerebral and cognitive dimensions of analysis. Our findings can be summarized at two levels: brain and cognitive.

At a brain level, we show that aging is associated with a large-scale reorganization of the language connectome based on simultaneous reduced functional specialization, increased integration, and deactivation of several subnetworks. These changes enhance the overall efficiency of language processing while minimizing the brain’s energy expenditure. At a neurocognitive level, we show that LP can be characterized as an inter-cognitive function influenced by the dynamic interaction between domain-general and language-specific (i.e., semantic) processes. Furthermore, our findings unveil that the emergence of LP decline during midlife may result from a decreased ability to reduce DMN activity. This reduction could impact older adults’ ability to retrieve semantic representations in a goal-directed manner, leading to difficulties suppressing irrelevant semantic associations during LP. Accordingly, our findings can be formalized as a novel model titled SENECA (Synergistic, Economical, Nonlinear, Emergent, Cognitive Aging), integrating connectomic (SE) and cognitive (CA) dimensions within a complex system perspective (NE) (Hancock et al., 2022).

From a connectomic perspective, our results align with previous work (Roger, Rodrigues De Almeida, et al., 2022), suggesting that language processes at rest depend on a large network composed of associative (RS NET1), sensorimotor (RS NET2), bottom-up attentional (RS NET3), and top-down control-executive (RS NET4) subnetworks. Within a lifespan perspective, we replicate previous findings showing that reduced segregation, reflected by a reduction of provincial nodes (Guimerà & Amaral, 2005), in control-executive, associative, and sensorimotor subnetworks (M. Y. Chan et al., 2014; Grady et al., 2016; R. Pedersen et al., 2021), can be considered a hallmark of aging, with a critical inflection point in midlife. Thus, this study confirms that midlife is a pivotal period for brain functional reorganization of the language connectome (Irwin et al., 2018; Lachman, 2015; Park & Festini, 2016).

Our results shed light on the neural mechanisms underlying lifespan functional changes. Specifically, we found that as individuals age, dedifferentiation, involving the over-recruitment of brain regions and reduced specialization (Fornito et al., 2015; Park et al., 2004), is consistently associated with enhanced functional integration and functional deactivation, which may constitute a compensatory strategy.

First, enhanced functional integration within the fronto-parietal (FPN) control network can be related to improved information transfer (Bagarinao et al., 2020), task processing flexibility (Bertolero et al., 2015; Tang et al., 2023), and overall better cognitive performance (Deery et al., 2023; Setton et al., 2021; Stanford et al., 2022). Through their precise adjustments of connectivity among adjacent regions (Bertolero et al., 2018), connector hubs imbue the network with integrative and flexible properties, thereby offsetting any decrease in specialization (as suggested by Cabeza et al., 2018). Consistent with this compensation account, our findings prove that the age-related integration of the FPN is not detrimental, as noted by Wu & Hoffman (2023). Instead, it serves as a beneficial mechanism, mitigating declines in executive functions during demanding cognitive tasks, particularly bolstering performance in multitasking, fluid intelligence, and language processing. Our study supports the idea that recruiting additional neural resources may represent a scaffolding response, actively fostering resilience in language processing throughout the lifespan (Park & Reuter-Lorenz, 2009; Reuter-Lorenz & Park, 2023). This also emphasizes that network dedifferentiation and compensatory integration are interrelated across the lifespan (Deery et al., 2023; Stumme et al., 2020).

Secondly, as individuals age, the increase in functional efficiency provided by integration appears to be closely intertwined with deactivation – the functional withdrawal of specific brain areas, including the left paracentral lobule, fusiform, posterior cingulate area (pCC), insula, and right IFG of the default-mode network (DMN; Alves et al., 2019; Menon, 2023). This deactivation or suppression of the DMN is a label given to the accumulation of peripheral nodes and has been well-established as an indicator of externally oriented attention, supporting demanding tasks by suppressing internal distractions like mind-wandering (Anticevic et al., 2012) and, more broadly, reducing task-irrelevant processes (Buckner & DiNicola, 2019). Specifically, the PCC has been emphasized as a hub between the DMN and FPN (Leech & Sharp, 2014), especially in tasks requiring controlled semantic access (Krieger-Redwood et al., 2016). Similarly, the right IFG has been linked with studies on age-related semantic fluency (Martin, Williams, et al., 2022; Meinzer et al., 2009, 2012). Thus, DMN suppression could mitigate age-related decline in goal-directed behavior, especially the semantic retrieval processes necessary for LP.

Our findings indicate the synergistic relationship between DMN suppression and FPN integration in the aging brain, reflecting a trade-off between functional efficiency and reorganization cost (Barabási et al., 2023; Barbey, 2018). This aligns with the brain’s wiring economy principle (Achard & Bullmore, 2007; Bullmore & Sporns, 2012; Shine & Poldrack, 2018), aiming to maximize the benefits of compensatory functional integration while minimizing energy expenditure through deactivation. In this context, our study proposed a probabilistic model to elucidate the functional mechanisms responsible for this “economic” reorganization throughout the lifespan, as illustrated in Figure 3.

Overall, our model shows a transition occurring in midlife within the language connectome, that is a shift from a more “energy-intensive” (costly) dynamic of compensation to a more “energy-efficient” one. Indeed, our research indicates that younger adults are more capable of accommodating the metabolic demands associated with sustaining long-range neural connections (Li et al., 2023; Liang et al., 2013; Tomasi et al., 2013). In comparison, older adults seem to adopt a more “energy-efficient” approach, substituting the reallocation of long-range connections between subsystems with short-range connections within subsystems, thus lowering the metabolic demands needed to achieve compensatory integration. This joins previous evidence showing reduced functional connectivity of long-range connections in older adults (Sala-Llonch et al., 2014) and may offer a metabolic explanation for the onset of cognitive decline observed in midlife in most cognitively demanding tasks (e.g., see Ceballos et al., 2024 for a study on the costs of brain dynamics).

At a brain network level, this shift is driven by a reduced synergy between DMN suppression and FPN integration as shown in Figure 4B. Indeed, compared to older adults, we found that younger adults capitalize on the cooperation or synergy between the DMN and FPN to enhance cognitive efficiency (Luppi et al., 2024; Xia et al., 2022). This is consistent with studies suggesting that the communication between higher-order networks on the sensorimotor-associative hierarchy, such as the DMN and FPN (Margulies et al., 2016), is more efficient at handling the high metabolic demands associated with long-range or rich club connectivity (Ceballos et al., 2024; Roy et al., 2017). Relatedly, our results are also consistent with the DECHA model (Spreng & Turner, 2019), proposing that reduced DMN suppression may result in a more inflexible modulation of the connectivity between the DMN and FPN in response to task challenges, *a fortiori* mediating cognitive decline in older adults, especially in lexical production.

While previous studies agree that DMN suppression is a vital element for regulating task-relevant dynamics (Leonards et al., 2023), as reflected by an increased metabolic response during tasks (Stiernman et al., 2021), the link between reduced DMN suppression and inflexible DMN-FPN coupling in older adults remains unclear. In this context, our study may bring elements of response by suggesting that reduced DMN suppression may raise the metabolic costs of maintaining long-range connectivity in the brain and that the inflexible DMN-FPN coupling is the consequence of older adults dealing with this increased cost.

Supporting this, previous research showed that flexibly allocating attentional resources from DMN to control regions, especially to the dlPFC as observed in our study (i.e., BA45 and inferior frontal sulcus areas), promotes fluid-related performances (Lu et al., 2022) maintaining a state of “global energy homeostasis” (Ramchandran et al., 2019), that is offsetting the cost of FPN integration as observed in our study. Consequently, reduced DMN suppression in older adults could raise the cost of FPN integration, momentarily disrupting this homeostasis. In response, older adults may thus prioritize allocating attentional resources through low-cost/short-range connections to restore the brain’s homeostatic balance, further supporting the shift from an “energy-costly” to a more “energy-efficient” integration around midlife. Interestingly, Ramchandran et al. (2019) also noted that homeostasis may be secured through “local cost-efficiency trade-offs,” thus being consistent with the discrepancy between a declining local efficiency but a preserved global efficiency across the lifespan (Cao et al., 2014; Song et al., 2014).

From a neurocognitive perspective, our results offer specific insights into the brain functional dynamics that uphold inter-cognitive functioning, specifically considering lexical production (LP), verbal fluency, and semantic abstraction. The main message is that the synergistic relationship between DMN suppression and FPN integration, characterized by flexible allocation of attentional resources via long-range connections, is a compensatory strategy that upholds inter-cognitive functioning in the aging brain. This compensatory dynamic aligns with a recent study suggesting that a youth-like network architecture that successfully balances integrative and segregation properties offers core resilience within the DMN and FPN networks (Stanford et al., 2022), translating to better semantic word-retrieval abilities (Krieger-Redwood et al., 2016; Martin, Williams, et al., 2022).

Importantly, our results show that LP (mainly lexical generation) is subserved by two distinct components, one Domain-General (DG) and one Language-Specific (LS), as posited by the LARA model (Lexical Access and Retrieval in Aging; Baciu et al., 2021). This confirms that LP can be viewed as an inter-cognitive function and the product of intra-(LS) and extra-(DG) linguistic processes (Gordon et al., 2018; Roger, Banjac, et al., 2022; Roger, Rodrigues De Almeida, et al., 2022).

On the one hand, the DG component, underpinned by multitasking, LP, and fluid-related abilities, was anti-correlated with DMN suppression and FPN integration across the lifespan. At a cognitive level, this aligns well with evidence linking the speed of retrieval and the generation of lexical predictions to fluid processing abilities (Brothers et al., 2017; Strijkers et al., 2011), further confirming that LP is a demanding task. At a neurocognitive level, this shows that reduced DMN suppression in older adults may compromise the ability to allocate resources from DMN to FPN regions cost-effectively, as discussed previously.

On the other hand, the LS component, underpinned by semantic performances and DMN suppression, peaked around midlife before declining in late life. While this confirms that individuals tend to accumulate semantic knowledge over their lives (Salthouse, 2019), this highlights that reduced DMN suppression in older adults also negatively impacts semantic abstraction performances, which is the ability to generalize semantic knowledge for efficient prediction (Moran et al., 2014), with implications for LP. Specifically, the increase in semantic performances in LS was proportional to the decline in fluid abilities. Given the constraint fluid processing places on LP, such accrual of semantic knowledge from younger to middle adulthood could represent a compensatory “semantic strategy” that maintains LP performance as fluid processing declines (Baciu et al., 2021; Wu & Hoffman, 2023).

Crucially, our results indicate that DMN suppression may capture the interplay between DG and LS as it covaries with both components. A reduction of this interplay could reflect how reduced DMN suppression in older adults compromises the ability to retrieve semantic knowledge in a goal-directed manner, manifesting at a cognitive level the difficulties in managing the cost of flexibly allocating attentional resources through long-range connections. Said differently, DMN suppression could index the controlled search and retrieval of semantic knowledge necessary for LP (i.e., DG-LS interplay) (Krieger-Redwood et al., 2019; Martin, Saur, et al., 2022). This is consistent with the notion that semantic cognition depends on both representational and control neural systems (Hoffman & MacPherson, 2022; Wu & Hoffman, 2023), with top-down processes regulating access to semantic representations (i.e., the DG-LS interplay; see also the Controlled Semantic Cognition framework proposed by Ralph et al. (2017).

As mentioned earlier, we suggested that the decrease in DMN suppression signifies a transition from a more “energy-costly” state to an “energy-efficient” one to maintain a global homeostatic balance within the brain. This transition aligns well with the lower control demands usually associated with the higher prevalence of “semanticized” cognition in older adults (Spreng & Turner, 2021). This also aligns with intriguing results revealing that older adults up until age 70 could continue to allocate (low-level) attentional resources, further delaying LP difficulties.

Indeed, after reduced DMN suppression in midlife, additional recruitment of the thalamus and the left putamen led to better semantic abstraction and LP performance (see Figure 4B). This is in line with studies showing that thalamic circuitry (Wolff et al., 2021; Wolff & Vann, 2019) facilitates functional interactions between multiple cortical networks (Badke D’Andrea et al., 2023; Gordon et al., 2022; Greene et al., 2020; Hwang et al., 2017), specifically maintaining stable representations at the semantic-lexical interface (Crosson, 2021). Importantly, both the thalamus and putamen showed high spatial concordance with the cingulo-opercular network (CON), whose benefits for word recognition have been emphasized in prior work (Vaden et al., 2013). This benefit could be attributable to the distinct role FPN and CON regions play in cognitive control (Sestieri et al., 2014; Wallis et al., 2015): the former managing externally guided/top-down control, selecting sensory cues from the environment, and the latter managing memory-guided/bottom-up control, providing sufficient flexibility for comparing sensory inputs to long-term memory traces (R. M. Brown et al., 2022). Thus, a synergy between CON and DMN connectivity changes could contribute to the more “energy-efficient” dynamic observed in older adulthood (Dosenbach et al., 2024; Han et al., 2023).

### SENECA: a novel integrative and connectomic model

To synthesize, our study extends theoretical accounts on neurocognitive aging, revealing that (i) LP draws from domain-general and language-specific processes; and (ii) the synergistic coupling between DMN suppression and FPN integration upholds inter-cognitive functioning across the lifespan while minimizing the brain’s energy expenditure.

As Livneh (2023) suggests, we combine the connectomic and cognitive levels of analysis within a single integrative model: SENECA. The SE-NE-CA model articulates a connectomic (SE) and cognitive (CA) dimension, which embraces a complex system perspective (NE). We borrow the name from the *Seneca effect,* generally found in complex systems, and describe “a slow rise followed by an abrupt decline” (Bardi, 2017), thus reflecting the dynamic of inter-cognitive functioning across the lifespan.

Along the connectomic dimension (S-synergistic; E-economical), we show that the synergistic relationship between DMN suppression and FPN integration provides the efficiency and flexibility necessary to compensate for reduced specialization while remaining economical – preserving global energy homeostasis. Along the cognitive dimension (C-cognitive; A-aging), we show that the onset of LP difficulties in midlife stems from reduced semantic control – the ability to exert control on accumulated semantic knowledge in a goal-directed manner – which may translate into poorer filtering of irrelevant semantic associations (Badre & Wagner, 2007; Barba et al., 2010; Jefferies, 2013).

Unifying both dimensions (N-nonlinear; E-emergent), reduced DMN suppression compromises the ability to manage the cost of FPN integration, prompting older adults to adopt a more “energy-efficient” strategy that preserves homeostasis at the expense of inter-cognitive functioning. This emphasizes that midlife is the turning point of a nonlinear and emergent neurocognitive dynamic marked by (i) a shift towards an increasingly semantic cognition to meet executive function decline (Spreng & Turner, 2021), and (ii) a shift towards a less synergistic coupling between DMN and FPN regions (Spreng & Turner, 2019). Importantly, SENECA aligns with a recent framework for cognition, suggesting that integration can be considered synergistic or redundant (Luppi et al., 2024; Mediano et al., 2022). Synergistic integration may correspond to “integration-as-cooperation” between networks as observed up to midlife (i.e., “energy-costly,” flexible DMN-FPN coupling). Redundant integration may correspond to “integration-as-oneness” within each network as observed beyond midlife (i.e., “energy-efficient,” inflexible DMN-FPN coupling).

In terms of clinical perspective, the SENECA model provides insights into the potential use of age-related neural mechanisms as biomarkers in midlife for predicting late-life neurodegenerative pathologies. Two hypotheses may be proposed: (i) Pathological word-finding difficulties before midlife may indicate increased maintenance cost of long-range connections, making them vulnerable to damage, as seen in pathological aging (Crossley et al., 2014). (ii) Beyond midlife, these difficulties may reflect challenges in securing a more “energy-efficient” compensatory integration strategy, compromising the brain’s homeostatic balance.

### Limitations of the study

Our study has several limitations: *(A) Cross-sectional data & Inter-individual variability*: The neural mechanisms identified in this study rely on probabilities derived from cross-sectional data, limiting their ability to account for inter-individual variability, especially in older adulthood (Stumme et al., 2020). Longitudinal studies would therefore be more appropriate to capture all the regions potentially underlying each mechanism. Future research should also investigate age-invariant mechanisms associated with high cognitive reserve for a more graceful language processing decline as age advances (Brosnan et al., 2023; Oosterhuis et al., 2023; Wen & Dong, 2023; Wulff et al., 2022). Further studies should specify the factors associated with the recruitment of subcortical structures, like the bilateral thalamus and left putamen, in late adulthood. *(B) Higher-order interactions:* Current graph theory methods assume dyadic relationships capture functional connectivity patterns of interest, but complex cognitive functions, such as language processing, involve higher-order interactions (Gazzaniga et al., 2019; Giusti et al., 2016). Modeling these interactions should be considered given the robust statistical and topological evidence (Schneidman et al., 2006; Yu et al., 2011; Gardner et al., 2021; Giusti et al., 2015). While SENECA does not explicitly address higher-order interactions, its predictions align with theoretical and empirical works showing age-related changes in synergistic interactions and their impact on cognitively demanding tasks (Gatica et al., 2021; Luppi et al., 2022; Rosas et al., 2022; Varley et al., 2023). *(C) Density thresholding & Atlas selection:* Sophisticated techniques are needed to account for the diversity of neuroimaging datasets and reduce intra-individual variability while preserving neurobiologically meaningful edges (Jiang et al., 2023). Proportional thresholding may exclude weaker edges, warranting exploration of data-driven filtering schemes like OMST (Orthogonal Minimum Spanning Tree; Dimitriadis et al., 2017) in understanding how older adults balance cognitive efficiency and reorganization cost. The SENECA model of neurocognitive aging pertains to the language connectome, but further investigation with whole-brain atlases is crucial, considering the impact of parcellations on reproducibility (Jiang et al., 2023; Ran et al., 2020).

## 5. Conclusion

This study aimed to elucidate the brain’s functional mechanisms responsible for the variation observed in language-related tasks as individuals age. Our findings highlight that, compared to language comprehension, the maintenance of lexical production (LP) depends on the synergistic relationship between suppression within the Default Mode Network (DMN) and integration within the Fronto-Parietal Network (FPN) in the language connectome. This relationship extends to support inter-cognitive tasks that draw upon both domain-general and semantic processes. Crucially, we propose that midlife represents a critical neurocognitive juncture that signifies the onset of LP decline, as older adults gradually lose exert control over semantic representations. This transition could stem from reduced DMN suppression which compromises the ability to manage the cost of FPN integration, prompting older adults to adopt a more cost-efficient compensatory strategy that maintains global homeostasis at the expense of LP performances. In summary, we encapsulate these findings in a novel integrative and connectomic model called SENECA, which articulates both cerebral (S-synergistic E-economical) and cognitive (C-cognitive A-aging) dimensions within the framework of complex systems (N-nonlinear E-emergent).

## Data availability

Raw data can be made available by the authors upon request. Scripts for analysis and visualizations are available at: https://github.com/LPNC-LANG/SENECA

## Supporting information

Appendix S2. Supplementary results.

Appendix S3. Reports the composition of RS LANG and the probabilistic resting-state network labeling of each LANG region.

## Acknowledgments

This work has been supported by the ANR project ANR-15-IDEX-02. This project has received financial support from the CNRS through the MITI interdisciplinary programs. The Cambridge Centre for Ageing and Neuroscience (Cam-CAN) research was supported by the Biotechnology and Biological Sciences Research Council (grant number BB/H008217/1).

## Conflict of interest statement

The authors declare no conflicts of interest.

**Appendix S1.** Supplementary material.

**Appendix S2.** Supplementary results.

**Appendix S3.** Reports the composition of RS LANG and the probabilistic resting-state network labeling of each LANG region.

