## Appendix S1. Supplementary material. for "Modeling the Neurocognitive Dynamics of Language across the Lifespan"

**MR Acquisition**

Resting-state functional MR images were acquired on a 3T Siemens TIM Trio with a 32-channel head coil for all participants at the Medical Research Council and Brain Sciences Unit in Cambridge, UK (MRC-CBSU). Two hundred and sixty-one volumes (261) were acquired with eyes closed in descending order using a gradient echo-planar imaging sequence lasting 8 min and 40 s (GEEPI, 32 axial slices, 3.7 mm thickness and interslice gap of 20% for whole brain coverage including cerebellum, TR = 1.97 ms, TE = 30 ms; voxel-size 3 x 3 x 4.44 mm, flip angle = 78°, field of view = 192 x 192 mm). In addition, structural images were acquired using a 1mm3 isotropic, T1-weighted Magnetization Prepared RApid Gradient Echo (MPRAGE) sequence and a 1mm3 isotropic (more information about the MR acquisition and resting state protocol are provided by Cam-CAN et al., 2014).

**Data Preprocessing**

The preprocessing of rs-fMRI data was performed using SPM12 (Welcome Department of Imaging Neuroscience, UK, http://www.fil.ion.ucl.ac.uk/spm/) running under MATLAB R2020b (MathWorks Inc., Sherborn, MA, USA). All images were realigned to correct for head motion and time-corrected with the mean image as the reference slice. The T1-weighted anatomical volume was co-registered to the mean image created by the realignment procedure and spatially normalized to the MNI (Montreal Neurological Institute) space before applying a smoothing function with a 6 mm FWHM (Full Width at Half Maximum) Gaussian kernel.

Motion parameters from the realignment were evaluated with ART (Artifact Detection Tool; Massachusetts Institute of Technology, available at: <https://www.nitrc.org/projects/artifact_detect>) to detect outlying volumes. We used an interscan movement threshold of 3 mm in translation, 0.02 rad in rotation, and a global interscan signal intensity of 3 *SD* relative to the session mean. In the next step, spatially preprocessed volumes were implemented in the CONN Toolbox for the FC analyses (Functional Connectivity Toolbox; Massachusetts Institute of Technology, available at: <https://www.nitrc.org/projects/conn>; (Nieto-Castanon, 2020).

**Table S1. Neuropsychological tasks**

| *Cognitive Domain* | Task | Descriptive Statistics (N=613; Mean, SD) |
| --- | --- | --- |
| *Language* | Naming  (Production - NAM) | M = 0.78, SD = .085 |
|  | Sentence comprehension  (Comprehension - SEN) | M = 0.89, SD = .074 |
|  | Proverb  (Semantic - PROV) | M = 4.57, SD = 1.6 |
|  | Tip-of-the-tongue (Retrieval - TOT) | M = -0.46, SD = .24 |
|  | Verbal Fluency  (Synergy - VERB) | M = 20.69, SD = 5.35 |
| *Domain-general* | Fluid Intelligence (FLUID) | M = 31.91, SD = 6.78 |
|  | Multitasking (MULTI) | M = -304.9, SD = 173.7 |
| *Long-term Memory* | Story recall (STRC) | M = 13.09, SD = 4.16 |

**Description of the neuropsychological tasks**

**NAM:** Name the pictured object presented alone (baseline), then when preceded by a prime object that is phonologically related (one, two initial phonemes), semantically related (low, high relatedness), or unrelated (Clarke et al., 2013).

**SEN:** Judge grammatical acceptability of partial auditory sentences, which begin with an ambiguous sentence stem (e.g., “Tom noticed that landing planes…”) followed by a disambiguating continuation word (e.g., “are”) in a different voice. Ambiguity is either semantic or syntactic, with empirically determined dominant and subordinate interpretations (Rodd et al., 2010).

**PROV:** Read and interpret three English proverbs (Huppert et al., 1994).

**TOT:** Participants are asked to name famous faces and indicate if they know/don’t know/or have a ToT (Brown & McNeill, 1966). In this study, we subtracted the score from 1 to stay consistent with a decrease as age increases (i.e., 1-x).

**VERB:** Mean of letter (phonemic) fluency and animal (semantic) fluency task. For the phonemic fluency task, participants have 1 min to generate as many words as possible beginning with the letter ‘p’. For the semantic fluency task, participants have 1 min to generate as many words as possible in the category “animals” (Lezak et al., 2012).

**FLUID:** Cattell Culture Fair Test: nonverbal puzzles involving series completion, classification, matrices, and conditions(Cattell & Cattell, 1960).

**MULTI:** Simulated tasks of a hotel manager: write customer bills, sort money, proofread adverts, sort playing cards, alphabetize a list of names. Total time must be allocated equally between tasks; there is not enough time to complete any one task (Shallice & Burgess, 1991). In this study, we subtracted the log from 1 to stay consistent with a decrease as age increases (i.e., 1-log(x)).

**STRC:** Listen to a short story, recall freely immediately after, then again after a delay, and finally answer recognition memory questions (Tulsky et al., 2003). Delayed recall measure used here.
