## Appendix S2. Supplementary results. for "Modeling the Neurocognitive Dynamics of Language across the Lifespan"

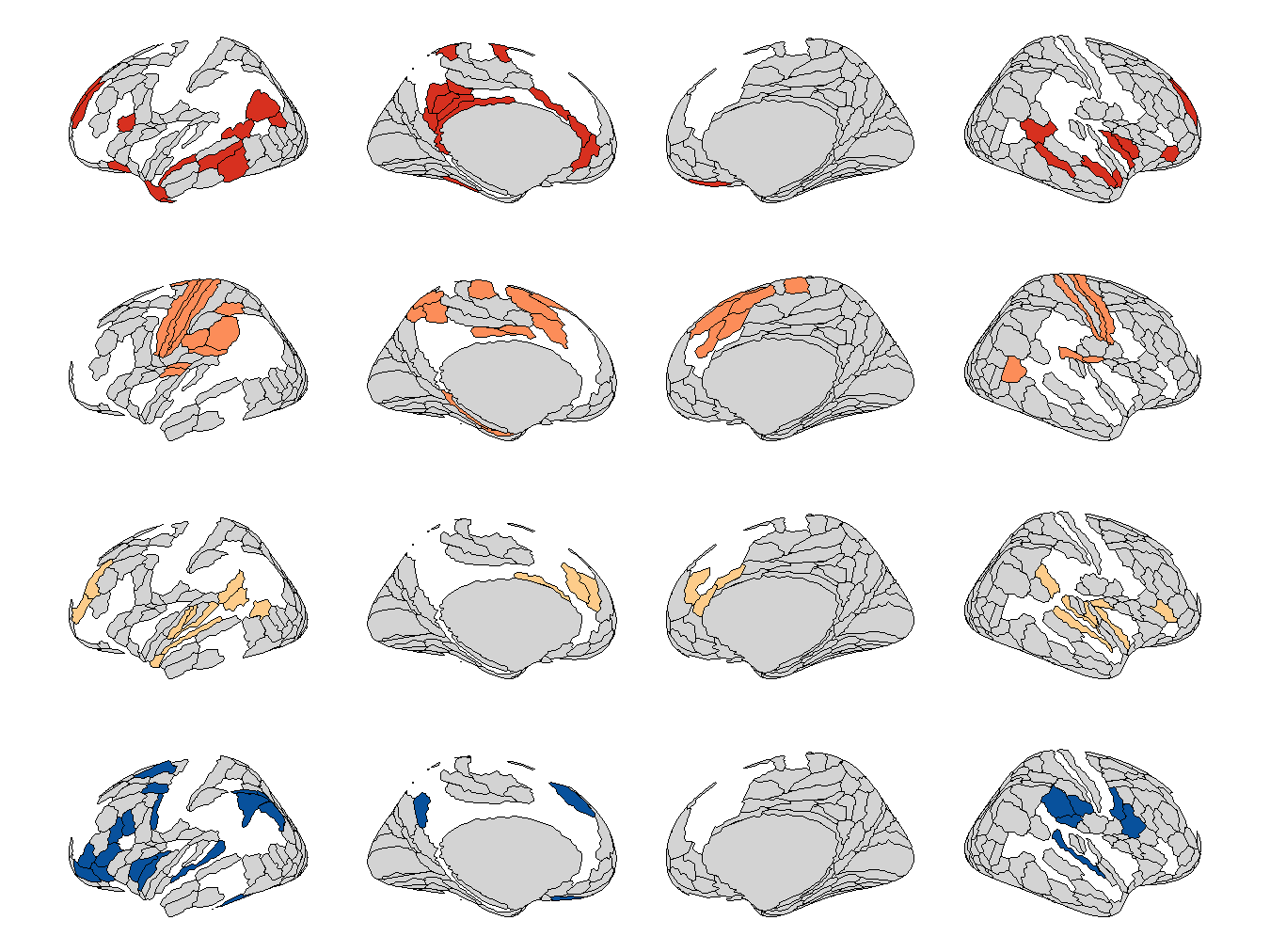

**Figure S1.** **Illustration of the rs-LANG connectomic atlas projected on the left and right hemispheres.** Projected on a multimodal parcellation of cortical (HCP_MMP1.0; Glasser et al., 2016). RS NETs: RS NET1 (40 regions, red), RS NET2 (34 regions, orange), RS NET3 (32 regions, yellow), RS NET4 (22 regions, blue). Each RS NET is a unique composition of resting-state networks (Ji et al., 2019). An additional 3-region module corresponding to the VMM (Ventral Multi-Modal) has been identified in the consensus partition but not considered for analysis due to its small size and low reliability across the lifespan. Brain visualization was created with the package *ggseg* in R (Mowinckel & Vidal-Piñeiro, 2020).

***Main differences with the task-based LANG organization***

We make three observations regarding the task-based counterpart (Roger et al., 2022) of the language connectome: (1) Whereas the language network forms its module to meet task-specific demands, all 11 regions at rest coalesce with the most extensive associative (45%) and bottom-up attentional subsystems (36%) while the remaining 20% forms crucial short-range connections with the control-executive subsystem. **This further suggests that core language processing at rest is functionally clustered with subsystems that support semantic/episodic access, and cognitive control.** (2) a third of CON (Cingulo-Opercular Network) regions are redistributed across the sensorimotor and control-executive subsystems. It suggests that regions supporting bottom-up attentional processes at rest may be more likely to tune their connectivity patterns with regions supporting sensorimotor and executive functions. (3) Finally, the consensus rs-LANG partition entails a 3-region module associated with the VMM (Ventral Multi-Modal) network, which tends to merge with the associative subsystem across the lifespan. Overall, this examination clarifies the critical differences in modular reorganization between intrinsic and extrinsic states of the LANG connectome (Figure S2). However, in-depth analysis may require a reassessment of the task-based LANG organization using our probabilistic approach for RSN assignment, particularly regarding the unlabeled subcortical structures in Roger et al. (2022).

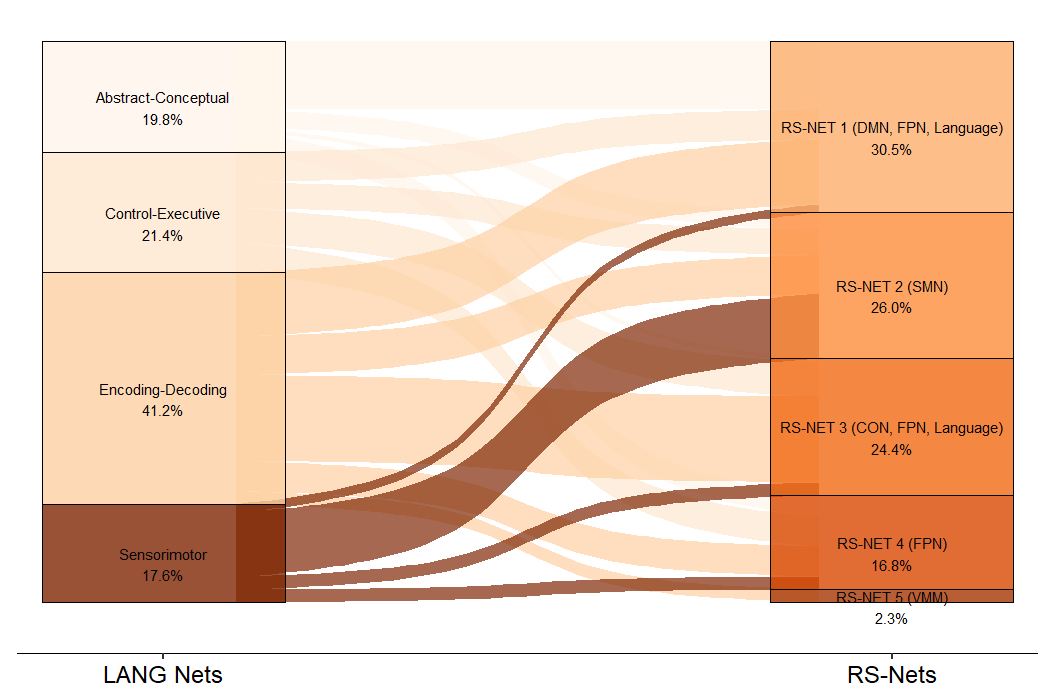

**Figure S2.** **Alluvial plot of the reconfiguration of the 131 LANG ROIs between the modular organization of the task-based LANG and resting-state-LANG.** Percentages correspond to the relative count of regions within each module relative to the whole LANG connectome.

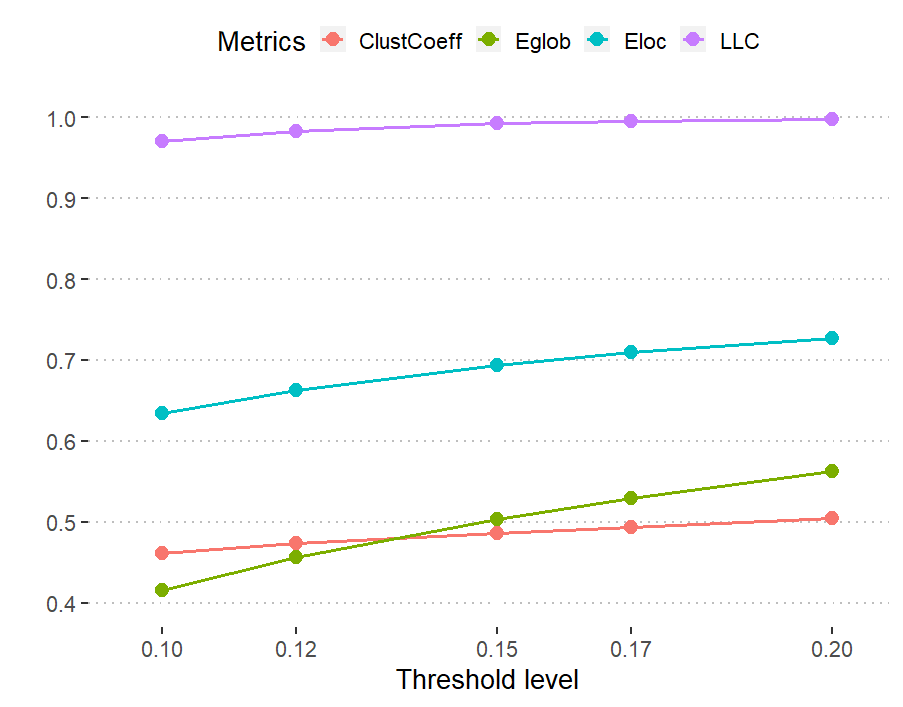

**Figure S3.** **Evolution of global metrics as a function of the threshold applied to the language connectome at rest.** The global metrics were calculated or averaged at the network level using the Brain Connectivity Toolbox (BCT; available at: <https://www.nitrc.org/projects/bct/>; (Bullmore & Sporns, 2009) implemented in MATLAB 2020b. Clust_coeff_: clustering coefficient calculated at the network level; E_glob_: global efficiency; E_loc_: local efficiency calculated at the network level; LLC: Largest connected component. 15% seems to be the optimal threshold at which nodes form tight clusters while preserving a relatively high global efficiency at the network level, denoting modular yet integrated processing.

**Subsystem-level analysis: validation of the consensus partition**

To ensure the consensus partition is well-fitted to all subjects, we repeated the same modularity analysis for younger, middle-aged, and older adults (18-39, *N* = 167; 40-59, *N* = 201; >59, *N* = 260). We used the AMI (Adjusted Mutual Information) to assess the similarity between the consensus and age group partitions. Analysis revealed that similarity was high for all age groups (AMI = 0.78, 0.91, and 0.83, respectively), suggesting that differences in the modular organization of rs-LANG are negligible across the lifespan.

**Small-world organization: Calculations**

We verified the small wordness assumption in the present study by computing the small worldness parameter. We estimated the mean path length and clustering coefficient we averaged over an ensemble of 10,000 Erdős-Rényi random graphs. At the selected sparsity level of 15%, we observed an average small wordness value of 2.31 [2.07, 2.51] CI 95%, thus verifying the assumption (small worldness above 1). The analysis script is available at <https://github.com/LPNC-LANG/SENECA>

**Probabilistic analysis**

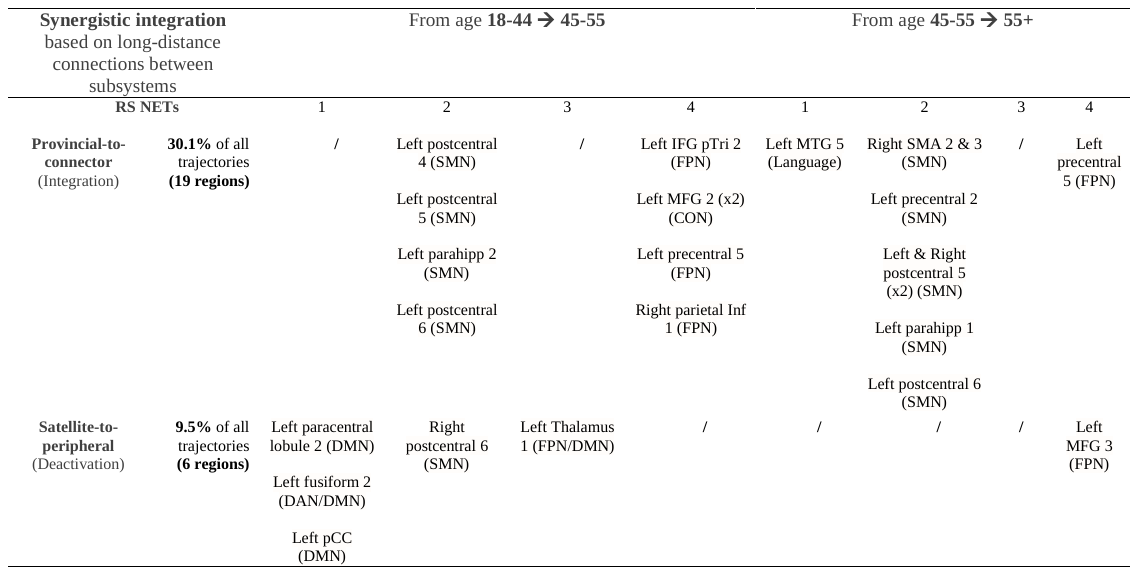

**Table S1 reports the regions that lose or gain long-distance connections across the lifespan**

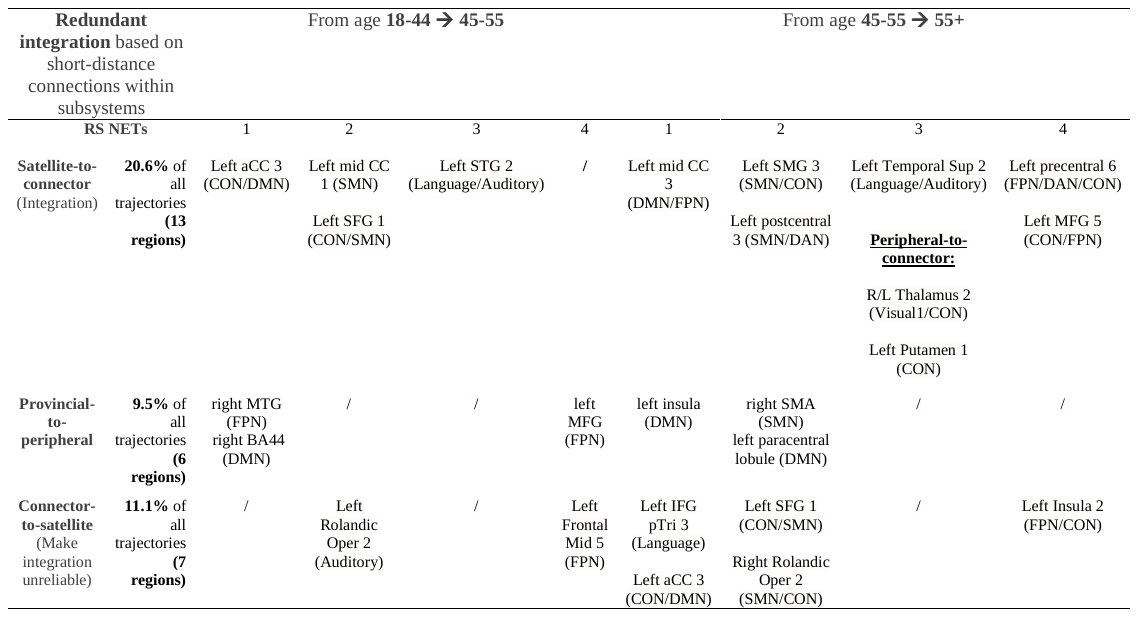

**Table S2 reports the regions that lose or gain short-distance connections across the lifespan**
